# Lira: Rotational Invariant Shape and Electrostatic Descriptors for Small Molecules and Protein Pockets based on Real Spherical Harmonics

**DOI:** 10.1101/2022.01.19.476747

**Authors:** Fernando R. Caires, Samuel R. Silva, Marcos Veríssimo-Alves, Vitor B. Pinheiro, Rinaldo W. Montalvão

## Abstract

**Motivation:** Modern AI-based tools are increasing the number of protein structures available, creating an opportunity and a challenge for automated high-throughput drug discovery pipelines. The amount of data is overwhelming for the current methods, thus demanding new high-performance approaches for Machine Learning-based rational drug design. As shape and electrostatics are the main components for understanding protein-ligand interaction; they are the primary targets for efficient AI-compatible descriptors and their associated comparison methods.

**Results:** The Lira toolbox is a set of components devised for describing, comparing and analysing shape and electrostatics for small ligands, peptides and protein pockets. It can generate databases with descriptors for tens of millions of shapes in a few hours, which can then be queried in seconds. The Lira design, focused on performance and reliability, makes its integration into AI-driven rational drug design pipelines simple.

**Availability and implementation:** Lira packages, available for download at https://pinheirolab.com/, are free to use for research and educational purposes.

## 1 Introduction

Small molecule compounds have dominated our pharmacopoeia from the very origins of the drug discovery field. Whether starting from natural compounds, close analogues or wholly synthetic leads, drug discovery, normally led by chemists, has built a long track record of success (Drews, 2000; Li and Vederas, 2009). The pace of success has been maintained at a high level through increasing investment and through the adoption of new technologies: be it increasing the throughput of candidates being tested or by improving the quality of data being used to guide the discovery process (e.g. using a high-quality structure of the protein target). With increasing frequency, computational approaches are being adopted to accelerate screening, design and automation of the process of drug discovery (Schneider, 2017), with machine learning and artificial intelligence methods the latest tool to the arsenal.

Drug discovery programs tend to focus their technology on the rapid assay of vast numbers of compounds. This random approach is called high-throughput screening and aims to identify compounds with IC50s lower than 10 *μ*M for their target proteins. This technology requires a considerable investment in faster systems for compound synthesis to generate large chemical libraries. Combinatorial chemistry using both solid-phase chemistry approaches and solution phase libraries coupled with high-throughput purification platforms is a requirement. Automation of bioassays and systems for collecting, storing, and analysing the extensive datasets generated is also compulsory. However, the rate of newly registered compounds in clinical trials has not increased in proportion to the exponential increase in investment brought about by these new robotic approaches.

Alternatively, it is possible to focus on targets and their related family members that are thought to be more tractable. The tractability of a target is based on the number of drug-like ligands for a target class and knowledge of the binding sites of family members using protein structure information. Classifying targets into families has allowed the design of focused compound libraries for particular families, as successfully demonstrated for protein kinases and various proteinases (Irving *et al*., 2001). Several approaches are now concentrating on screening tiny molecules, or ‘fragments’ from which a lead can be designed using knowledge derived from biophysical assays of how the fragment binds in the active site of the target (Li, 2020).

Lately, modern *in silico* approaches for identifying potential drug candidates are taking centre stage. Ligand docking aims to find the optimum binding position and orientation for a compound in the active site of the proteins, with the best docking programs correctly docking most ligands when tested on large sets of protein-ligand complexes. However, difficulties arise in predicting the affinities of the different compounds for the protein active site. Regardless, virtual screening is helpful in docking and ranking a large number of compounds so that the highest-ranking compounds can be selected for acquisition or synthesis and experimentally tested for activity against the target protein. Virtual screening significantly enriches for accurate hits in a selected subset of compounds (Bender and Glen, 2005).

The fragment-based approach and virtual screening are designed to provide a more reliable and effective sampling of chemical space, thus minimising the number of tested compounds and reducing the cost of drug development. Such optimisation of the small molecule screening process has allowed experimental methods such as NMR, X-ray crystallography and cryo-EM to be harnessed for drug discovery. Fragments are typically tiny organic molecules of between 100 and 250 Da and low binding affinities (below 10 mM) against the target proteins. Consequently, they cannot be identified using traditional high throughput screening, and biophysical methods must therefore be used to detect the fragments in the active site (Blundell *et al*., 2006). Once a functional fragment has been identified and its binding mode defined, it is incorporated into a template for a larger ligand with better drug qualities. Although the fragment hits have a low affinity, they frequently exhibit high ligand efficiency, i.e. high values for the average free energy of binding per heavy atom, and this property makes them strong candidates as starting-points for drug design and optimisation.

While fragment-based structure-determination-aided drug discovery is successful, it is a resources-heavy approach and potentially not sustainable for targets that are not easy to crystallize or whose structure cannot be readily determined. As computational costs drop and methods improve, an entirely *in silico* approach is feasible. With the advent of modern AI software for high-quality protein structure prediction, such as AlphaFold (Jumper *et al*., 2021) and RoseTTAFold (Baek *et al*., 2021), the possibility of replacing the experimental methods with pure in *silico* drug design pipelines is closer than ever.

Such virtual drug discovery pipelines can potentially reduce the waste of time and resources poured into research and development dead-ends. Unfortunately, current drug discovery pipelines rely heavily on human-based intervention for many specialised decisions. Therefore, a new generation of AI-based approaches for virtual drug discovery and design is needed since the complexity and amount of data involved in such decisions is overwhelming.

One crucial part of any drug discovery pipeline is the method used for ligand similarity analysis. One of the classical methods for shape-only similarity is USR (Ballester and Richards, 2007), a high-performance inter-atomic metric. Although the molecular shape is an important descriptor, it alone is insufficient for constructing a robust ligand dissimilarity metric. The USRCAT (Schreyer and Blundell, 2012) is a USR extension that adds pharmacophoric information to deal with the shape descriptor limitation without significant performance penalties. Other methods, like ElectroShape (Armstrong *et al*., 2010) and eSim (Cleves *et al*., 2019), rely on information about the molecular electrostatic potential to enrich their descriptors. Even though those methods are moving in the right direction, by adding electrostatic information, they employ very simplistic calculations compared to Quantum Mechanics approaches. In addition, they do not provide a unified framework for ligand superposition and comparison.

We designed Lira to fulfil the ideas proposed by Morris *et al*. (2005) about the use of Spherical Harmonics as 3D pocket and ligand shape descriptors. Although critical, many of the proposed concepts were beyond the technology of the time. Notably, Density Functional Theory or other Quantum Mechanics methods for computing the surface electrostatic potential were prohibitive due to their computational cost. With the advance of affordable and powerful hardware, many methods for high-performance computing have become available.

Our approach builds on the pioneering ideas of those methods by creating rotational invariant and comprehensive descriptors for surface shape and electrostatic potential. In particular, our descriptors are based on a Neural Network (Rathi *et al*., 2020) that reproduces Quantum Mechanics methods level-of-quality for the computation of the surface electrostatic potential. Compared to the previous methods, our approach has the benefits of a high-performance and controllable coarse-to-fine representation designed specifically with Machine and Deep Learning applications in mind. In particular, we are interested in the potential of combining Deep Generative Models, like DeLinker (Imrie *et al*., 2020), with Lira to build the initial leads from fragments and use Lira for database screening, thus creating an automated high-throughput *in silico* drug discovery pipeline.

## 2 Approach

The use of Spherical Harmonics as descriptors for mathematical functions defined on the Unit Sphere is a unifying view of their geometric properties, as it not only includes a complete representation but provides the structure for their rotation (Green, 2003). Such geometric representation can approximate the shapes and electrostatic potentials of molecular surfaces or protein pockets, with the additional advantage of being compact and easy to compare. Our approach takes advantage of these properties to construct sets of well-defined geometric descriptors ready to be applied on Machine Learning or Deep Learning methods involved in rational drug design.

## 3 Methods

Here, we propose a framework for comparing surface shapes and electrostatic potential through three distinctive layers of increasing complexity. All layers share the same mathematical foundation, constructed around the three-dimensional representation of star-shaped functions by Spherical Harmonics.

### 3.1 Real Spherical Harmonics

Spherical harmonics (SH) are an important class of orthogonal functions whose linear combinations can express any real square-integrable function of *θ* and *Φ* defined in the 3D space. The series expansion of a given function *f* (*θ*, *Φ*) in SH is

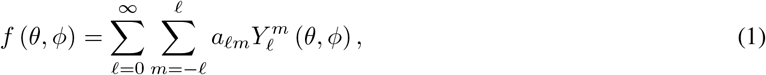

where *f* (*θ*, *Φ*) is the function to be expanded, 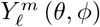 a particular SH, *a_lm_* its coefficient, *θ* is the polar angle, *Φ* is the azimuthal angle (ISO 80000-2:2019 convention), and *𝓁* and *m* are the SH degree and order, respectively.

In this work, we are interested in the particular case where the SH are defined as real values by:

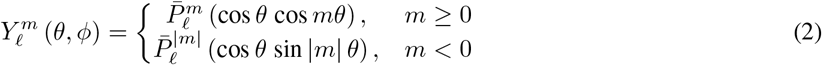

In equation (2), 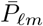 is the normalized associated Legendre polynomial, defined as:

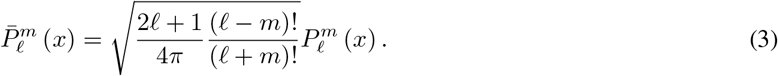

As a consequence of this normalization, any squared SH integrated over the sphere is unity:

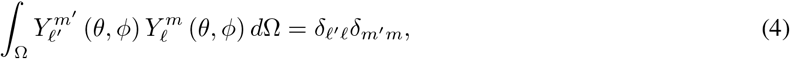

where *dΩ* = sin *θ dθ dΦ* is the differential surface area on the unit sphere. Multiplying both sides of equation (1) by 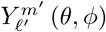, integrating over the sphere and using equation (4), it is straightforward to demonstrate that the *a_lm_* can be calculated as:

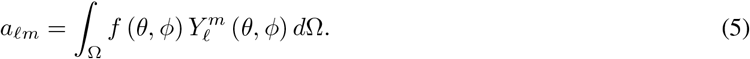

### 3.2 Molecular Surface Shape and Electrostatic Potential

In Lira, the function *f* (*θ*, *Φ*) can describe either the radius of a molecular surface or its electrostatic potential at that radius (fig. 1). Those functions are defined through a triangle mesh generated by the ESP-DNN program (Rathi *et al*., 2020) that contains a triangulation of the molecular surface and of the potential at each vertex position (fig. 2). The ESP-DNN program generates an electrostatic potential (ESP) surface through a graph convolutional deep neural network (DNN) model trained on DFT-generated ESP surfaces for 105,500 molecule models filtered from a set of 1,336,480 commercially available ones, using the method succinctly described below.

**Figure 1:**
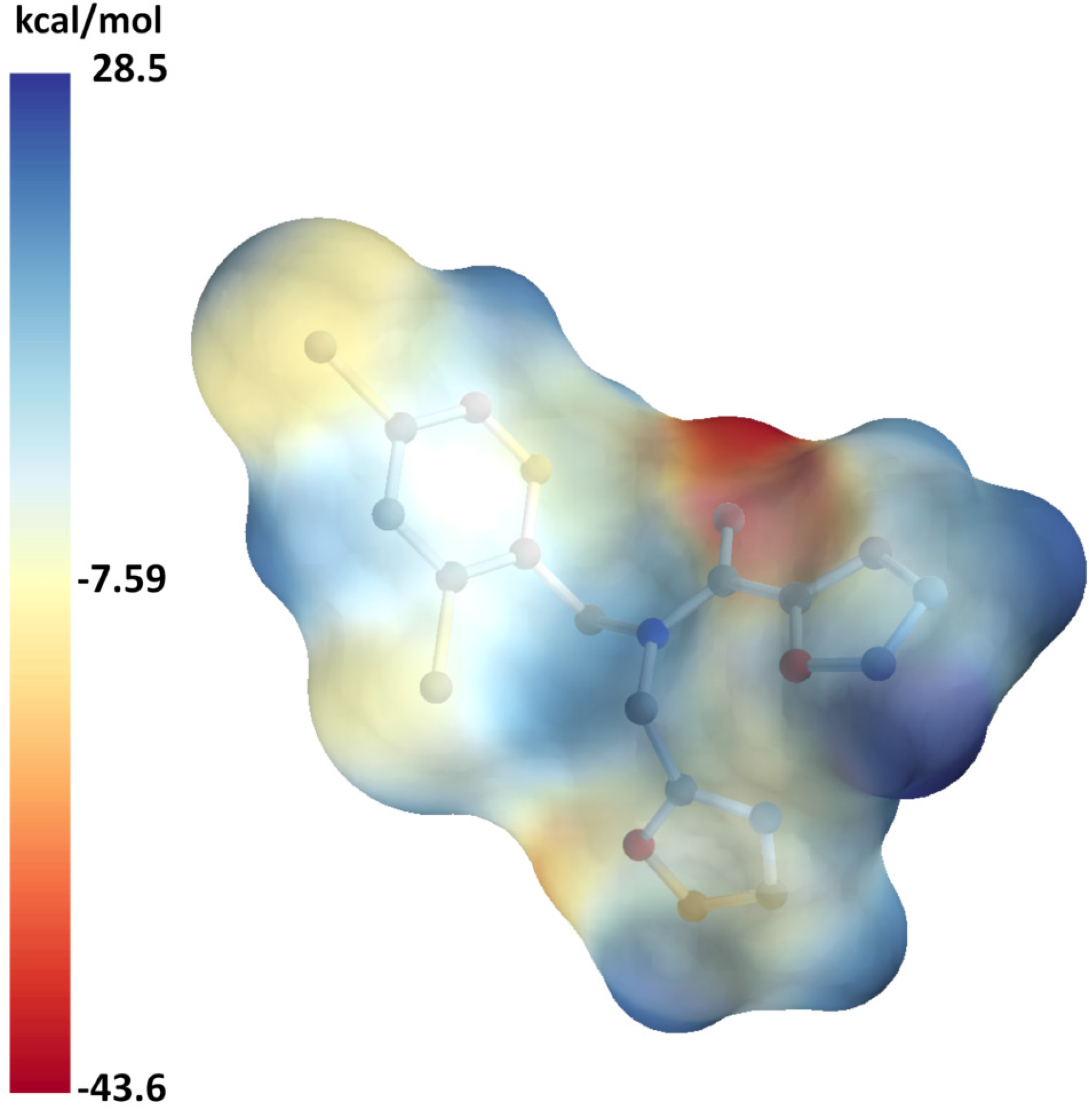
Example of a molecular surface generated by the ESP-DNN program using a Neural Network for emulating the Quantum Mechanics calculations. This molecule ZINC28231927 is used as a running example for all subsequent calculations and analyses.

**Figure 2:**
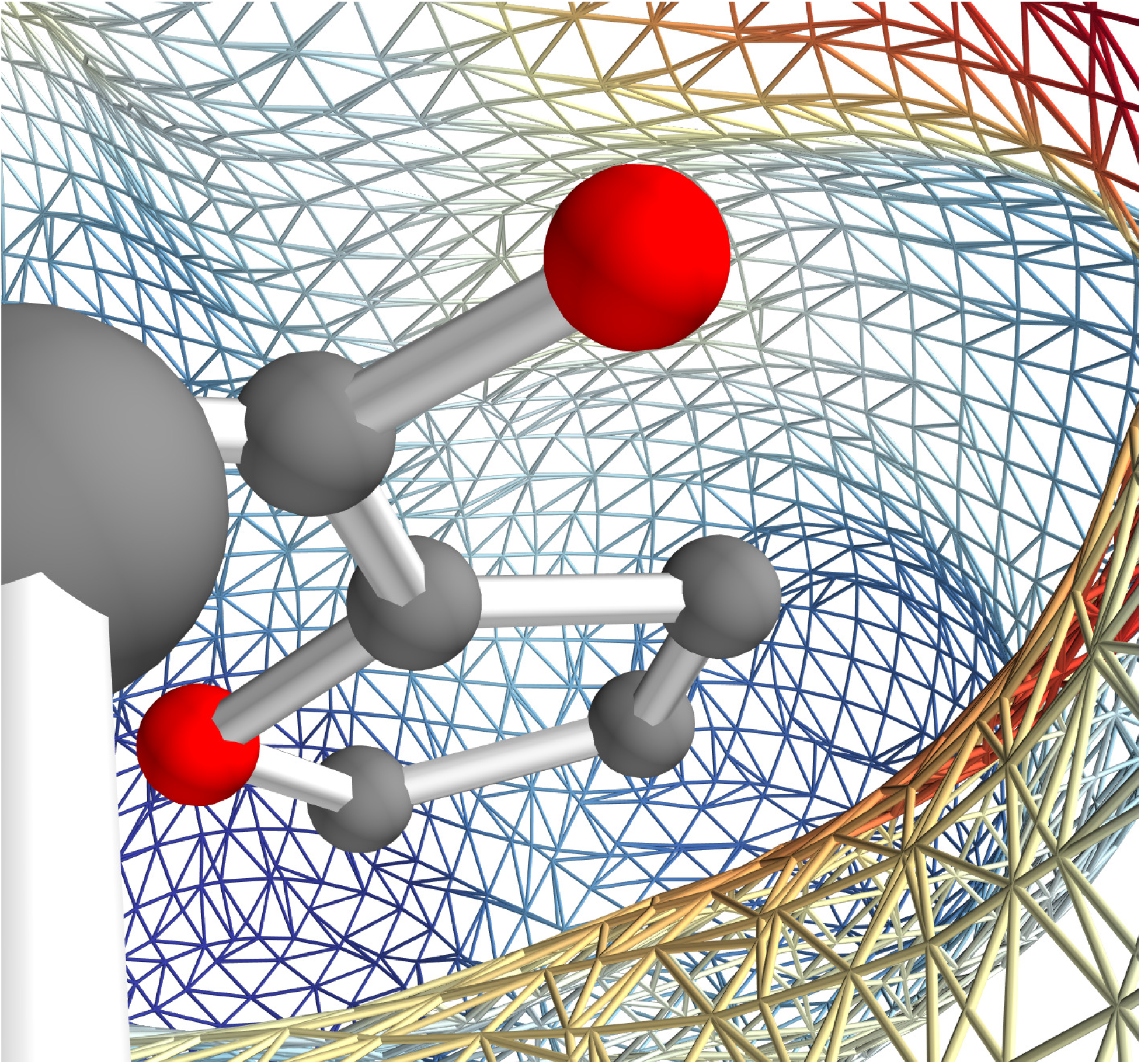
Interior of a molecular mesh generated by the ESP-DNN program.

First, feature groups containing non-hydrogen and hydrogen atoms of a molecule, plus lone pairs, p orbitals and *σ*-holes are generated, each of which having point charges associated to them, for all molecules of the set. Molecular geometries were optimized by DFT at the B3LYP/6-31G* level of theory and ESP surfaces are then generated by single-point calculations at the B3LYP/6-311G** level of theory. Next, the point charges for the feature groups are fitted to optimally reproduce the DFT ESP surfaces, generating the DFT-fp model for point charges. Finally, DFT-fp results are used to train a molecular graph convolutional DNN that will then generate values for the point charges associated to feature groups of molecules which do not belong to the training set (DNN-fp). The quality of the DNN-fp ESP surfaces is essentially as good as that of the DFT-fp, and much superior to the ESPs obtained by the AM1-BCC method, widely used in codes such as ANTECHAMBER. Moreover, the time for executing a DFT-fp calculation (~0.3 s) is about 100 (6,000) times faster than the corresponding AM1-BCC (DFT) calculation. This speed is crucial for the generation of ESPs for large numbers of molecules to be queried by Lira.

### 3.3 Monte Carlo Integration

As the mesh description for the surface and its electrostatic potential are not symbolic functions, it is necessary to evaluate equation (5) through Monte Carlo Integration (Green, 2003). Equation (5) can thus be rewritten as:

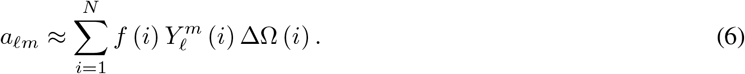

We are sampling *i* = 1… *N* points regularly distributed over the **unit sphere** and, in this case, the approximate element of surface area 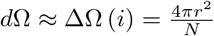 is just 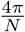. Since the sampling points i, randomly determined by the Monte Carlo method, are not necessarily the same as the mesh vertices, we employ the Möller–Trumbore intersection algorithm to find which triangle the associated ray intersects (Schlick and Subrenat, 1993). The intersection point, for very small triangles, is where the projected sampling point touches the molecular surface. The value for the electrostatic potential at this position can be evaluated by an interpolation of the 3D triangle using barycentric coordinates of the projected sampling point relative to the vertices in which electrostatic values are calculated (fig. 3).

**Figure 3:**
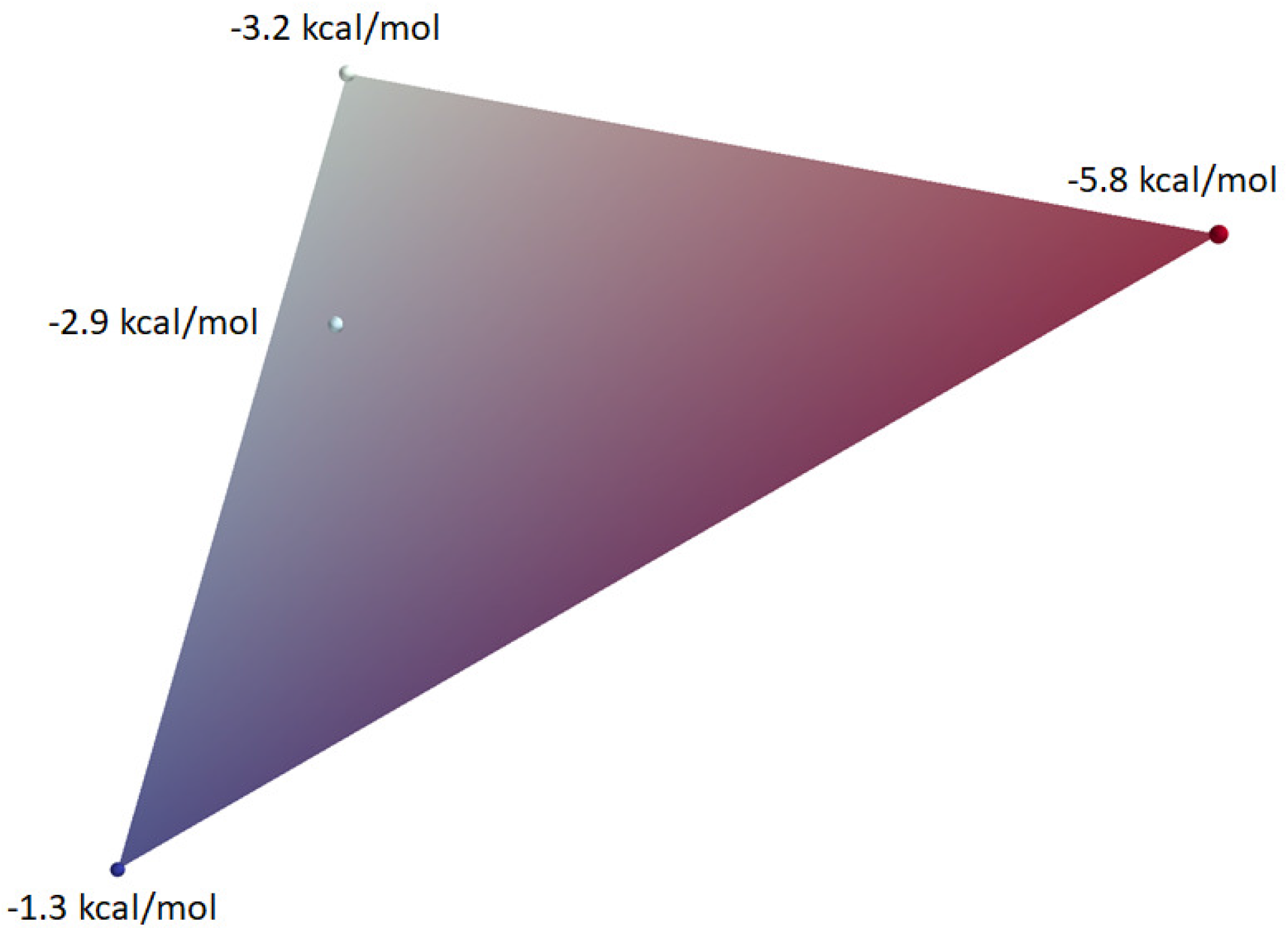
Möller–Trumbor projected intersection and barycentric interpolation of a single spherical sampling.

### 3.4 Energy Rotational Invariant Descriptor

Equation (1) allows the reconstruction of the surface and its electrostatic potential by truncating the expansion of *f* (*θ*, *Φ*) at a maximum value of the SH degree *𝓁_max_* (fig. 4). This surface shape and electrostatic representation have a coarse-to-fine nature; thus, their details increase as the maximum SH degree level increases. A carefully selected *𝓁_max_* creates a small number of SH coefficients that describe the dual shape-electrostatic information in a very compact form (fig. 5). By comparing the Pearson correlation coefficient between the originally sampled values and the reconstructed ones for 18,701,140 ligands, we selected *𝓁_max_ =* 16 as the reconstruction shows good agreement with the original values (figs. 6 and 7).

**Figure 4:**
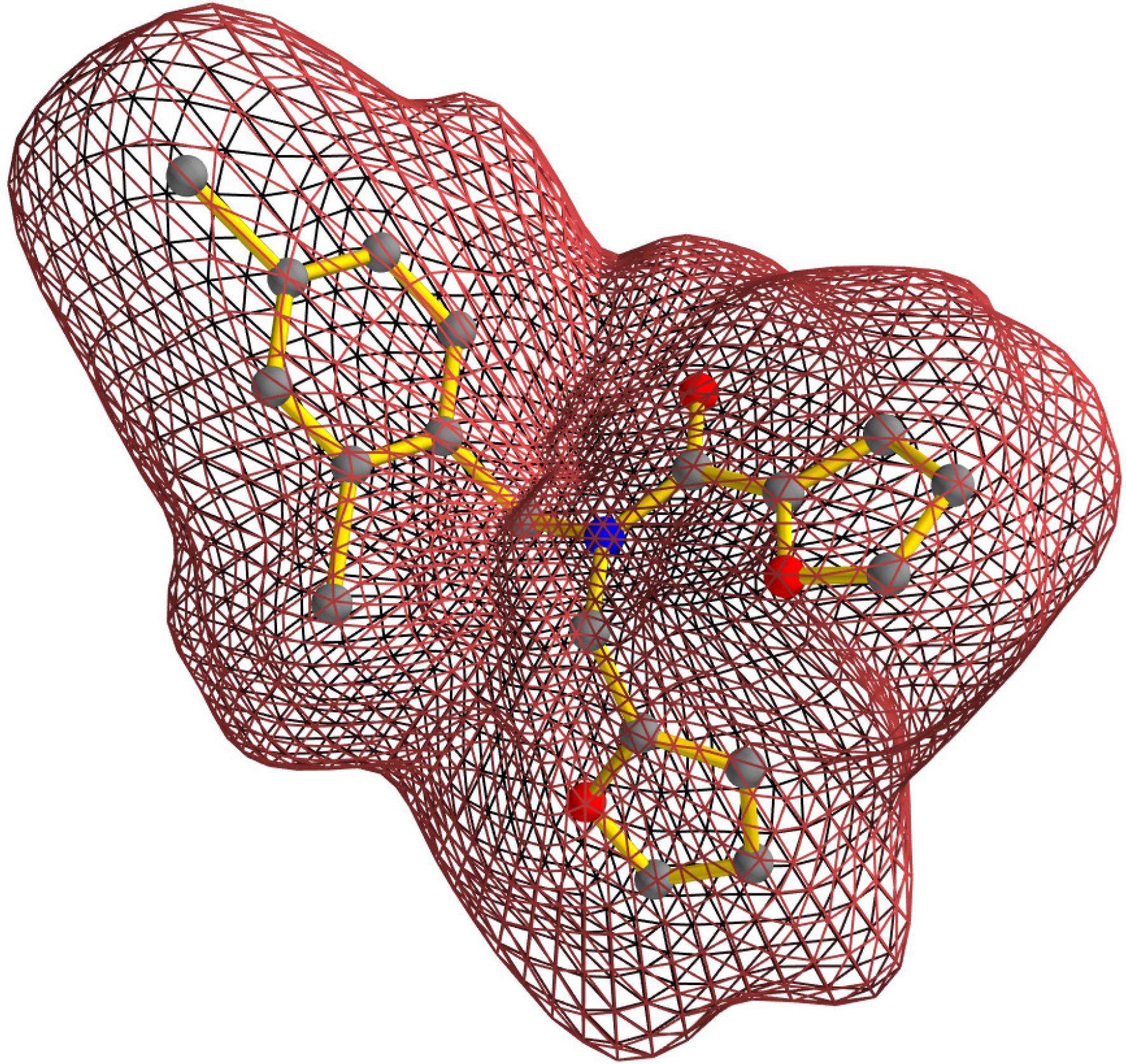
Molecular surface shape reconstruction using *𝓁_max_* = 16.

**Figure 5:**
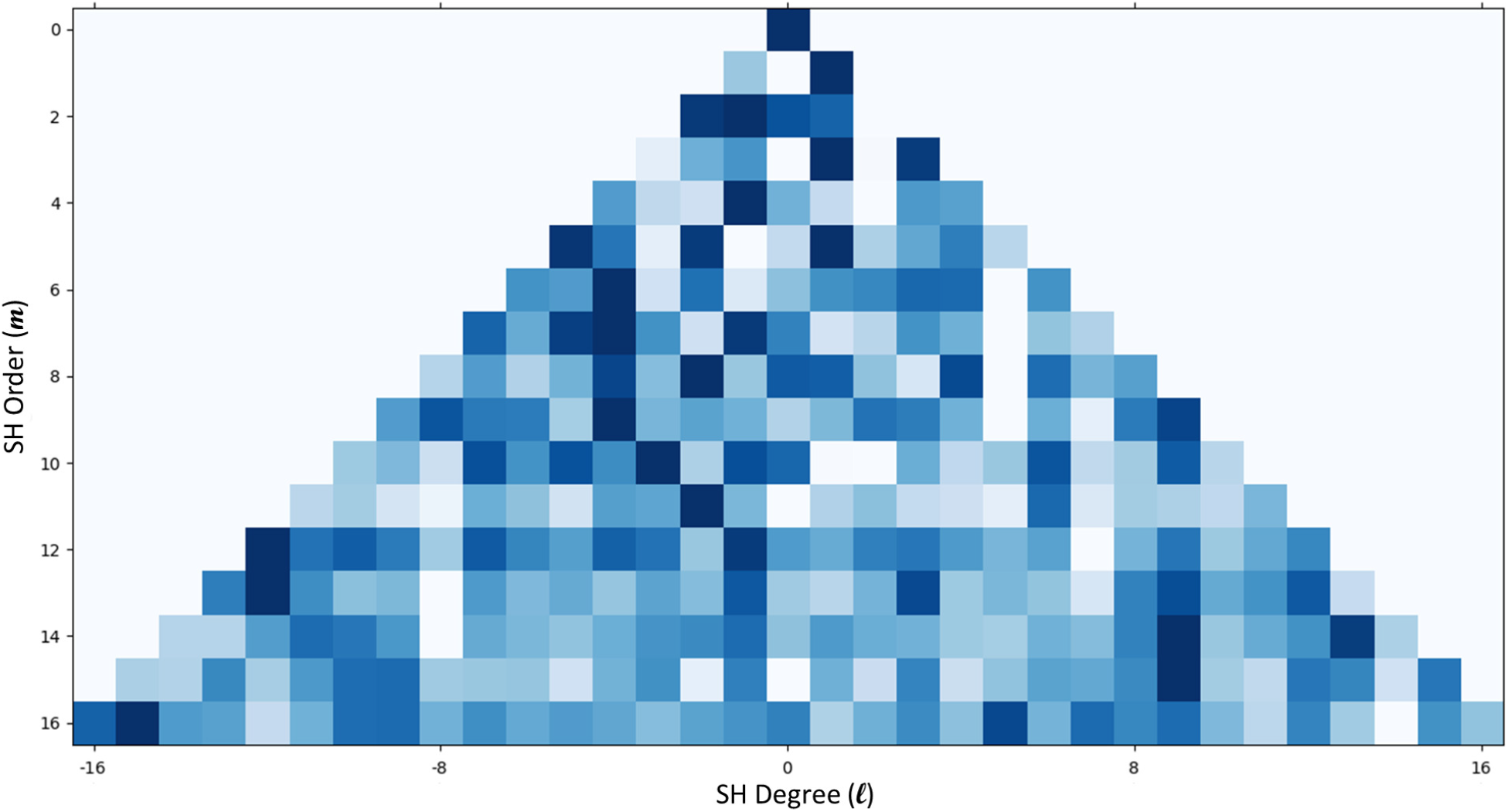
Molecular surface shape SH coefficients for *𝓁_max_* =. For illustration purposes, the coefficients values are normalized by row.

**Figure 6:**
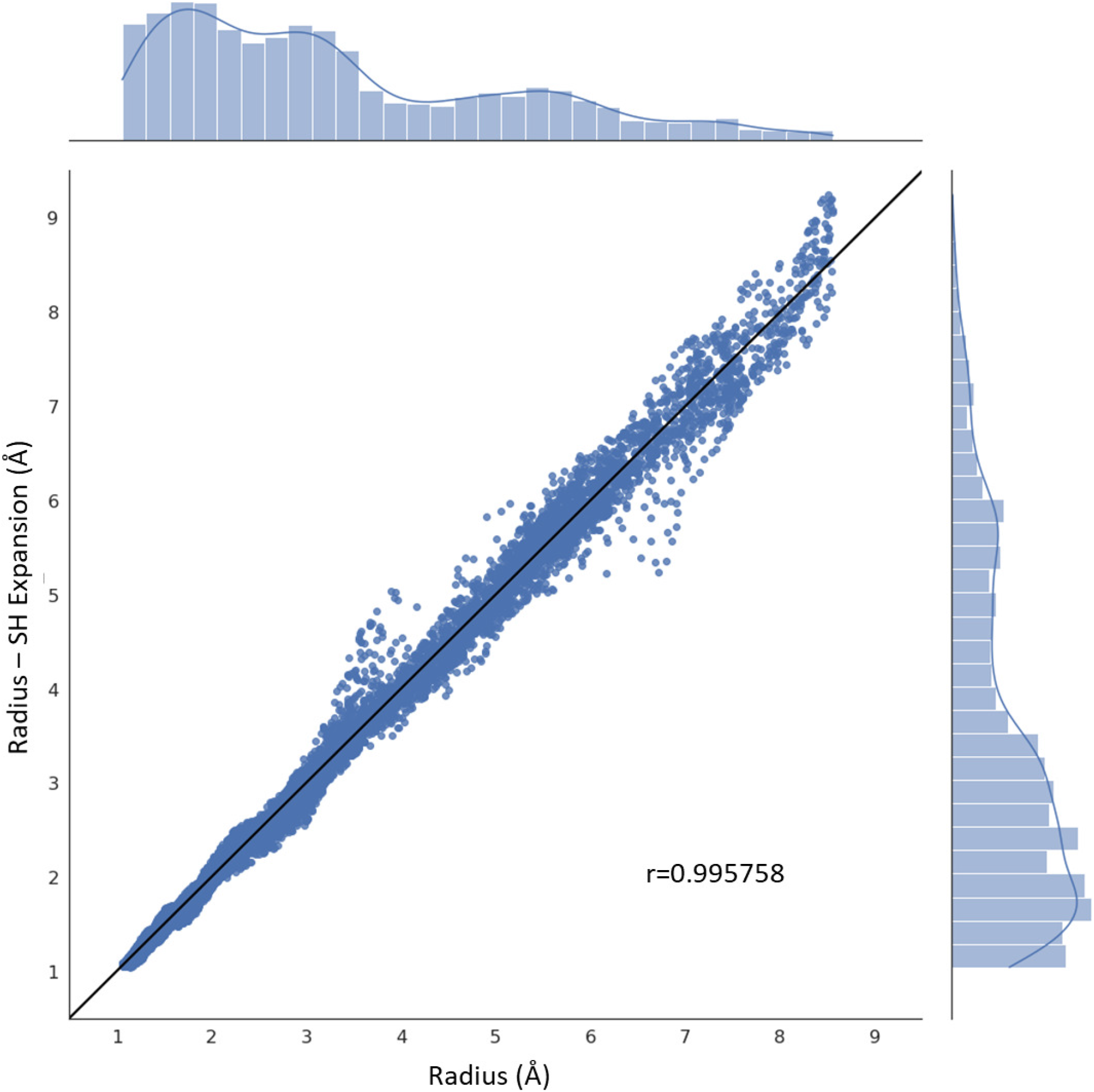
Comparison between original and reconstructed shaped radius for the surface of a ligand.

Two sets of Energy Rotational Invariant Descriptors (EI), one for the shape coefficients and another for the electrostatic ones, are constructed by adding all squared coefficients of each SH degree *𝓁* up to *𝓁_max_* = 16 in a variation of what has been suggested by Li and Hartley (2007):

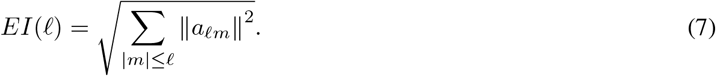

Equation (7) does not constitute a complete descriptor, thus suffering from ambiguity, and the errors produced by comparing two descriptors are mainly false positives. In other words, using this descriptor to measure ligand dissimilarity is likely to identify a set of ligands including structures that are not entirely similar. The idea behind the use of this descriptor for both shape and electrostatics is that they act as a first coarse-grained sieve, demanding minimal computational time and effort. This first sieve is capable of comparing and classifying tens of millions of ligands in under a minute, selecting a suitable number of compatible shapes and electrostatics for further high-quality analysis.

Figure 8 shows an example of such an EI computed for the SH coefficient set of the molecular shape shown in figure 4. The first EI (*𝓁* = 0) shows a much higher value than any other component. That is easy to understand if we look at how each SH function contributes towards the surface representation: Looking at figure 9, which shows the first seven real SH functions, we can see that *𝓁* = 0 corresponds to the spherical contribution for the shape expansion. As in this specific case the ligand shape is more spherical than elongated (fig. 4) it is to be expected that this particular SH would dominate the expansion.

**Figure 7:**
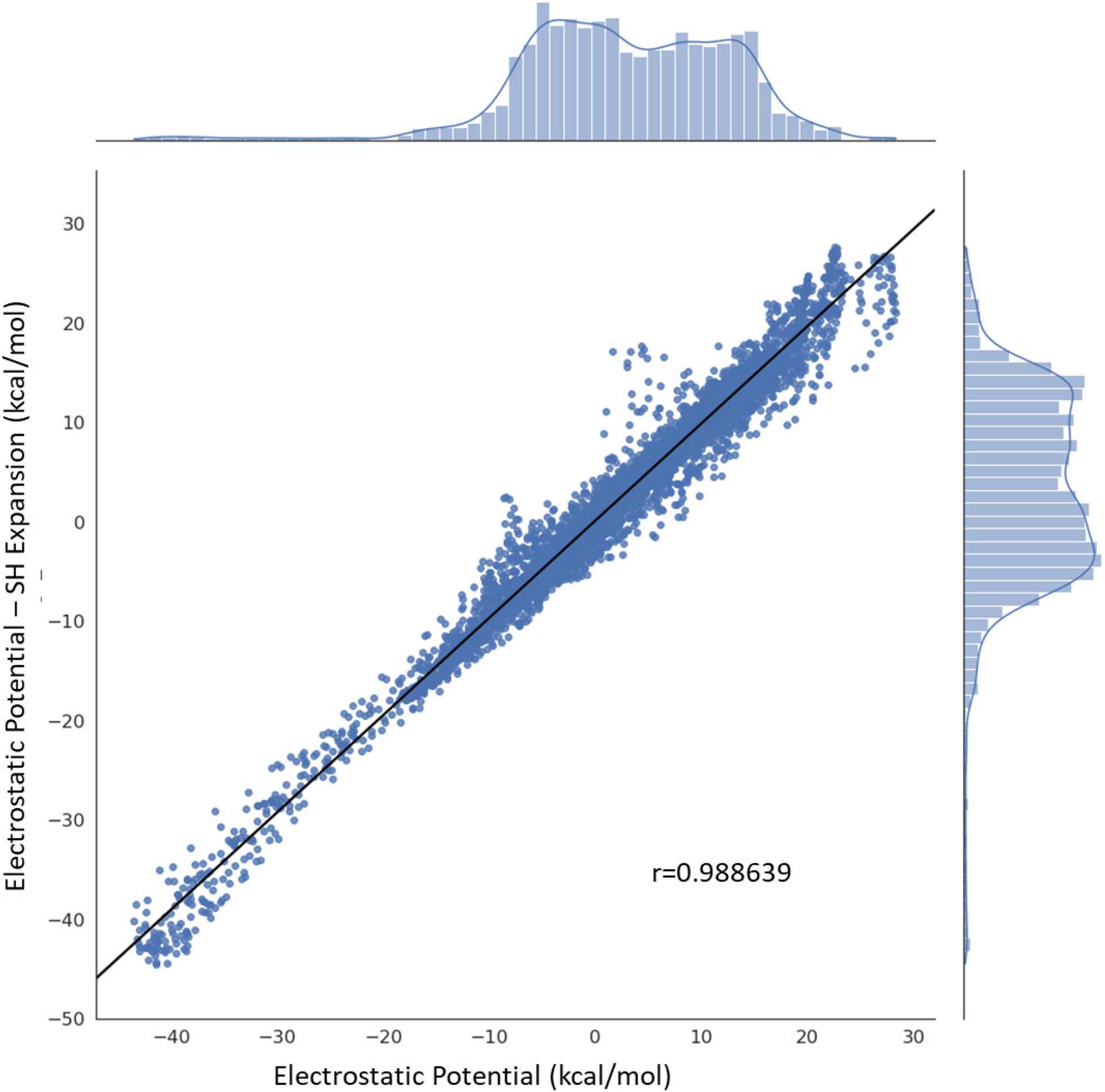
Comparison between original and reconstructed electrostatic potential at a ligand surface.

**Figure 8:**
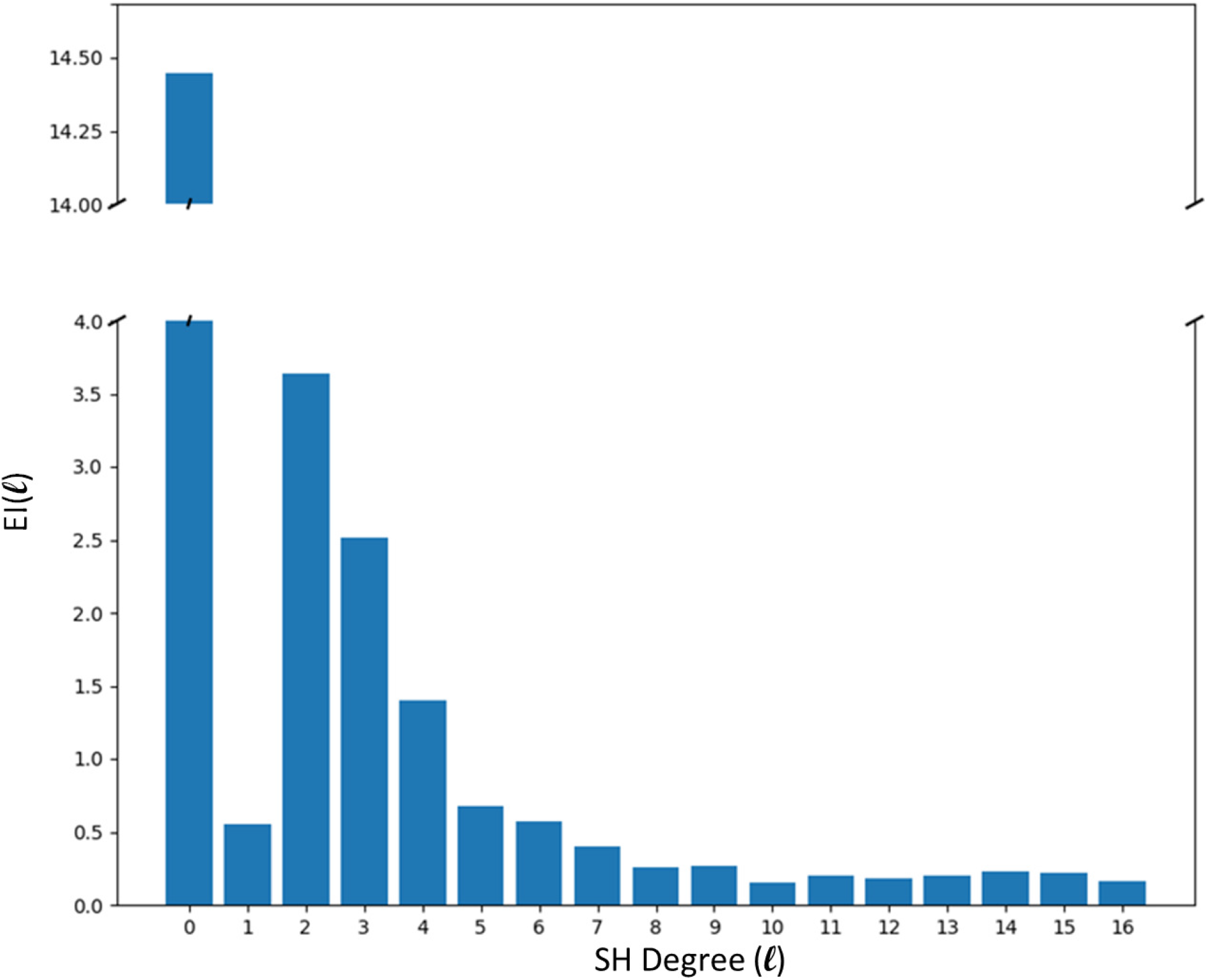
Molecular surface shape EI for *𝓁_max_* = 16.

**Figure 9:**
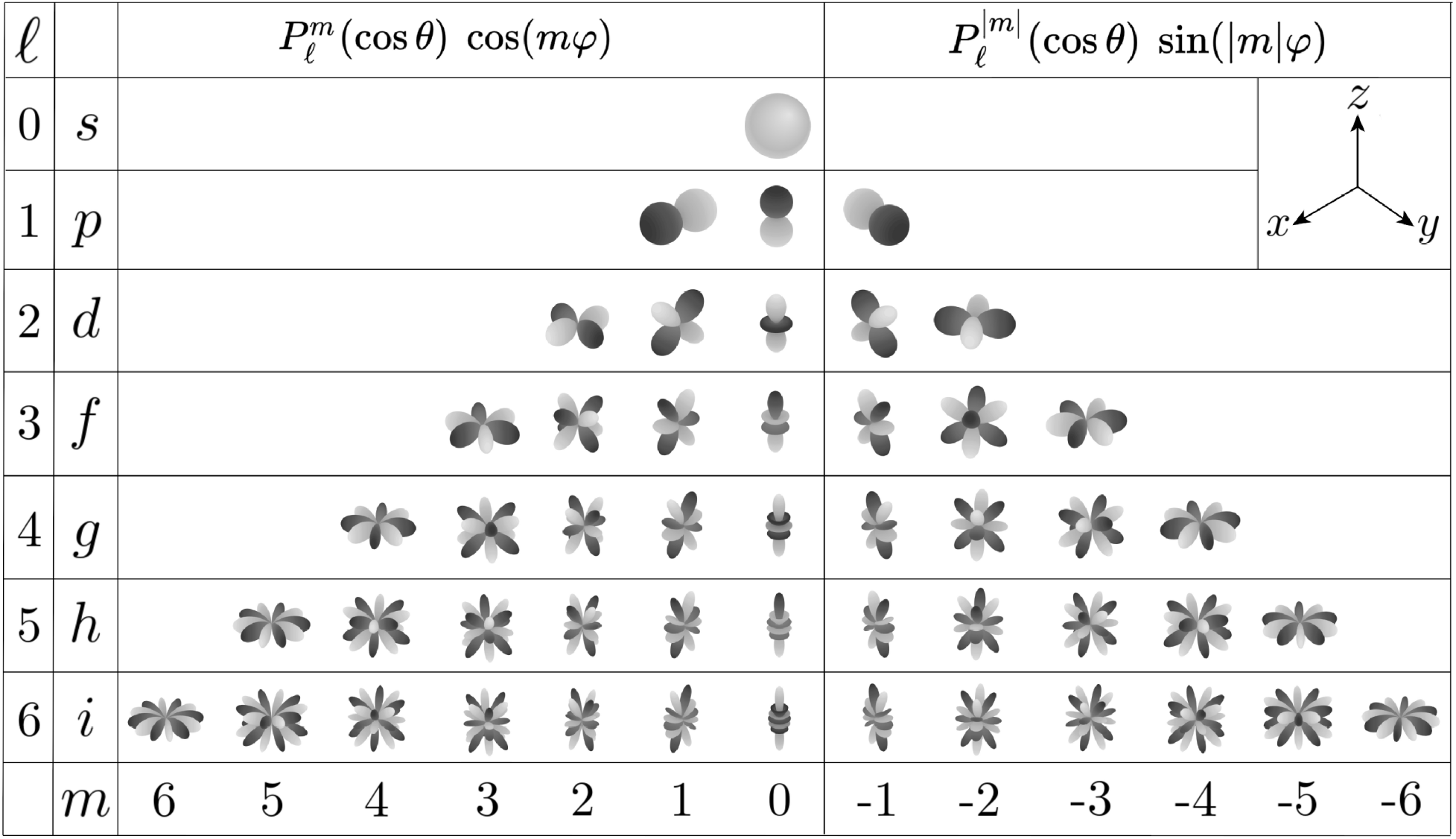
Examples of Real Spherical Harmonics functions 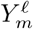. (Adapted from https://commons.wikimedia.org/wiki/File:Sphericalfunctions.svg)

### 3.5 Voxel Grid Rotational Invariant Descriptor

The second descriptor used in Lira is based on a rotation invariant representation of a voxel grid (Kazhdan *et al*., 2003). The Voxel Grid Rotational Invariant Descriptor comprises a spherical harmonic representation of concentric spheres of different radii. In order to compute this invariant, it is necessary to restrict the voxel grid of the shape to a collection of concentric spheres. The next step is to represent each spherical restriction in its SH expansion. The final step is the computation of the norm of each frequency component for each radius. This computation results in a rotational invariant 33-element vector representation of a 2D grid indexed by radius and frequency.

Although this representation only covers SH frequency components for shape, the use of concentric spheres allows the algorithm to better characterize the intricacies present in many ligand/pocket surfaces. This vector makes it possible to refine the comparison between two different shapes and remove false positives as result of steric clashes. This refinement is crucial when comparing all shapes present in a dynamical ensemble of a ligand, instead of just comparing one particular pose.

### 3.6 FFTs on the Rotation Group SO(3)

The final method in Lira allows us to compare all the (*𝓁_max_* + 1)^2^ = 289 SH coefficients for shape or electrostatics. As these SH coefficients are not rotationally invariant, a reproducible and efficient method is necessary to rotate them for aligning two shapes together. The alignment is achieved through the implementation of the ICP algorithm (Besl and McKay, 1992) by Funkhouser *et al*. (2004) using the SOFT 1.0 SH library (Kostelec and Rockmore, 2003). This technique determines the Wigner-D rotation matrix that aligns the two sets of SH coefficients by calculation of a FFT on SO(3). Given two surfaces with reconstructed functions *f* (*θ*, *Φ*) and *h*(*θ*, *Φ*), the correlation function *C*(*g*) between them is:

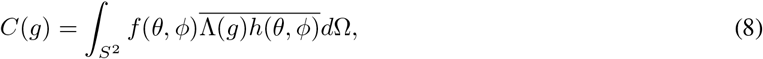

where *g* ∈ *SO*(3) and *Λ*(*g*) is the linear operator that produces a Wigner-D function when applied to 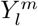. Essentially, the algorithm finds the *g* that maximizes the correlation function (eq. 8) between the two shapes described by *f* (*θ*, *Φ*) and *h*(*θ*, *Φ*) (eq. 1). By applying the optimal Wigner-D rotation matrix to the SH coefficients of one of the functions, the algorithm aligns both shapes into an orientation where the direct comparison of all coefficients is possible. Figure 10 summarizes schematically the three sieves that compose the Lira algorithm.

**Figure 10:**
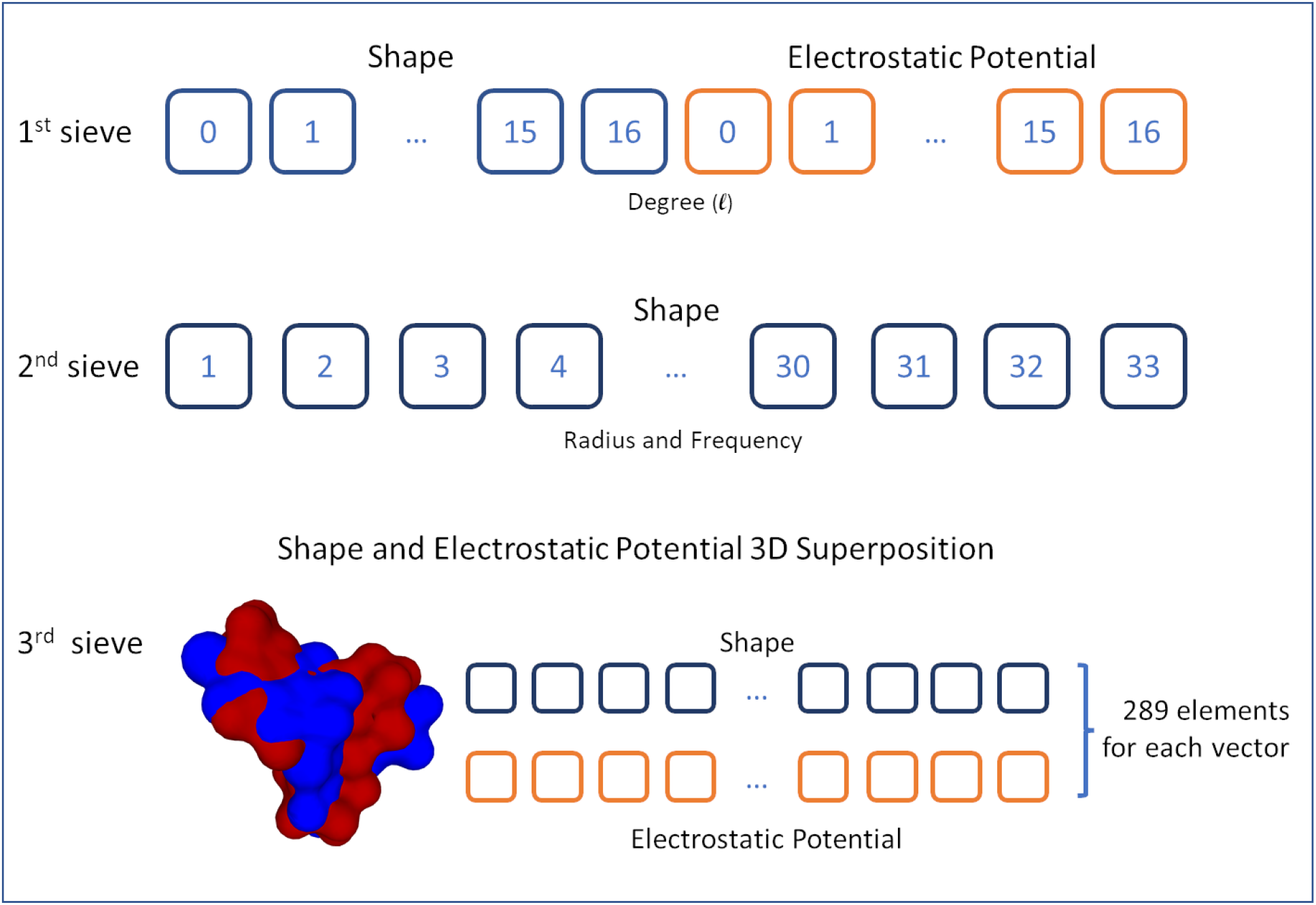
The three layers for surface shape and electrostatic potential comparison.

### 3.7 Descriptor Comparison

The final aspect of our method is the comparison between two descriptors. Using the procedures described in the previous sections, Lira produces three different sets of mathematical descriptors (fig. 10) for shape and electrostatic potential. Although slightly different, we can compare them using the same metric. As the scale for each *𝓁*-band is different, we use the weighted Euclidean distance, as its construction is very suitable for such cases:

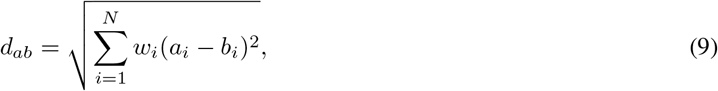

where *w_i_* can either be 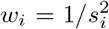 (the inverse of the *i*-th variance) or a user-defined measure of importance of the *i*-th element of the descriptor, *a_i_* and *b_i_* are elements of two descriptor vectors and *N* is the number of elements. The metric for 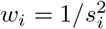 is a particular case of the Mahalanobis distance, known as standardised Euclidean distance. As the weight is the reciprocal of each variance, it effectively places all the measurements onto the same scale. It implies that the variations in each of the coefficients are equally critical, even though they are measured on a different scale.

## 4 Implementation and Discussion

Having established Lira as a robust platform for comparing molecules, we set out to demonstrate it in the ZINC database (Sterling and Irwin, 2015). While it is possible to carry out comparisons on the fly, it is more efficient to carry out these calculations once and use Lira to query this new database. We called this new locally-generated database Z4NC, since it adds 4D (3D shape + 1D electrostatic potential) to ZINC.

### 4.1 Database Preparation

The first step for creating the Z4NC database is storing each ligand MOL2 file in the RockDB storage using the filename as a key and its contents as the value. The RockDB is a persistent key-value store optimized for fast storage that allows the Z4NC database to retrieve any structure given its id.

For each file, we apply the ESP-DNN method to compute the surface triangle mesh with its electrostatic potential, and we compute their SH expansion. Due to the large file size of the triangle mesh, typically over 1MB, we chose to store it only during the SH coefficient computation phase. When stored in binary format, the coefficients are only 578 bytes (289 + 289 for *𝓁_max_* = 16) and more suitable for long-term storage.

The next step is to compute the two sets of coarse energy rotational invariant descriptors directly from the SH coefficients for the surface and electrostatic potential (eq. 7). The more refined voxel grid rotational invariant descriptor is computed only for the molecular shape by directly applying the Kazhdan *et al*. (2003) method to the triangle mesh.

The `lira_coeffs` program, a high-performance code in the D programming language, is responsible for computing the SH coefficients from the triangle mesh and the energy rotational invariant descriptors. The `ShapeDescriptor` C++ program, authored by Kazhdan, computes the voxel grid rotational invariant very efficiently. The total computational time in an AMD Ryzen Threadripper 3990X-based computer using 64 cores is around 230 hours, and it is done only once during the creation of the database. All the generated data is efficiently stored in a HDF5 (The HDF Group, 2010) file using the `PyTables` (PyTables Developers Team, 2021) library.

### 4.2 Database Search

Once created, it is possible to search the database for molecules of similar shape and electrostatics as one or more query molecules. To do so, one needs to compute the RIDs for the query molecule and use the `lira_search` program to find a set of matching molecules. The search procedure comprises three sieving phases. In the first phase, a collection of 50,000 molecules is selected from the initial set, in ascending order of dissimilarity of the energy rotational invariant descriptor. The next phase prunes this initial collection by using the voxel grid rotational invariant descriptor to select the top 500 molecules (ranked by dissimilarity).

The final molecule collection is small enough for shape alignment using the `ShapeAlign` C++ code by Kazhdan *et al*. (2003). It applies the *SO*(*3*) FFT to align all the shapes, thus giving us a dissimilarity measure for all the 289 SH coefficients using equation (9) and setting *w_i_* = 1. Although it is a representation that depends on the molecule orientation, the complete SH coefficient set is non-ambiguous and fully describes the surface and its electrostatic potential. Morris *et al*. (2005) extensively discuss the application of the complete SH coefficient set for aligned molecules using this same metric. Nevertheless, that method only employs shape descriptors, and their heuristic approach for shape alignment is not entirely deterministic or optimal (Morris *et al*., 2005). The *SO*(*3*) FFT is unique and optimal, thus producing an aligned reference framework for all selected molecules.

### 4.3 Search Results and Analysis

One of the essential applications of ligand dissimilarity analysis is the mapping of the molecular vicinity, as the typical query molecule is just a barely adequate fit for the target pocket. Using different original query molecules, we tested our method by searching the Z4NC database for the best matches and comparing the results with the original query molecule. Figure 11 shows the top six results, ordered by distance, to the query molecule ZINC43833531, shown in 1. All molecules selected by Lira show highly similar shapes and surface electrostatic potential. The whole search process takes less than one minute using one CPU core of our AMD Threadripper 3990X and the Julia language version of `lira_search`.

**Figure 11:**
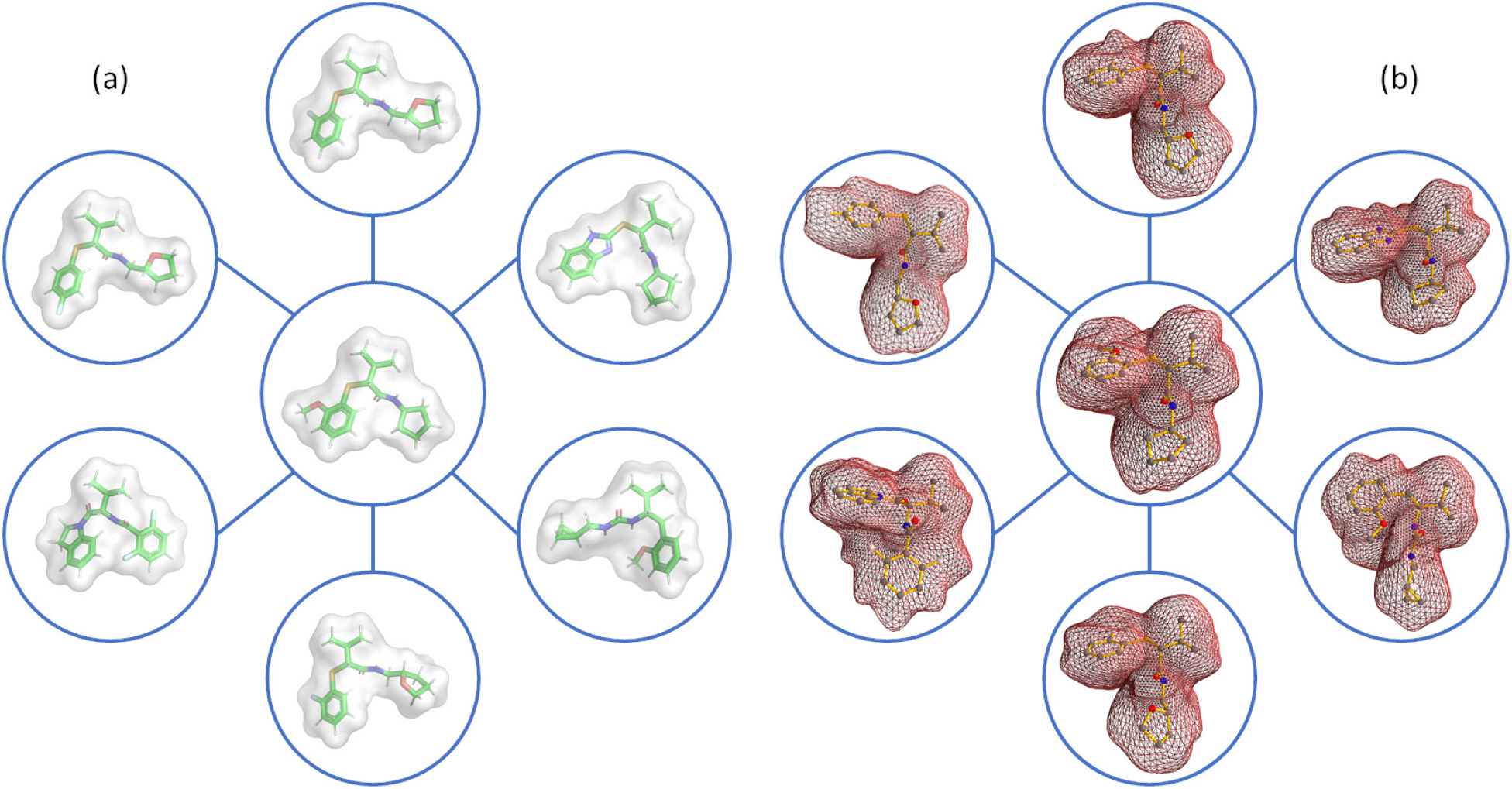
A comparison of some molecules selected by Lira based on the query molecule ZINC67007472 (shown in the centre of the diagrams). The best selected molecules clockwise from the top are: ZINC43833531, ZINC42901627, ZINC81475237, ZINC43833520, ZINC21278029 and ZINC43831578. (a) shows the original molecular surfaces and (b) the SH reconstructed ones for *𝓁_max_* = 16, although not in the same orientation as a for clarity.

As shown in Figure 11, SH distortions, due to the order cut-off value, can be observed. However, the distortions are systematic and thus do not significantly impact the molecular selection process (see Figures 6 and 7). The 10,000 points being used to sample the molecule surface remain accurate and explain the high quality of the initial selection.

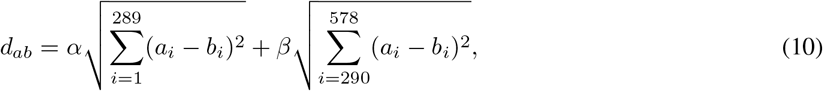

The first two sieves are fast and powerful enough to produce a large set of well-adjusted molecules in the vicinity of the query molecule. Although large, it is still small enough for more complex clustering algorithms like HBSCAN (McInnes *et al*., 2017) or for the third sieve to apply dendrogram exploration through Deep Learning. Equation 10 describes the distance matrix for the third sieve, where *α* and *β* are shape and electrostatic potential importance weights and *a_i_* and *b_i_* are elements of two descriptor vectors containing the SH coefficients. A Deep Learning algorithm can easily navigate the dendrogram and select molecules that emphasize the shape (figure 12c) or the electrostatic potential (figure 12d) compatibility with the query molecule (figures 12a and 12b) by adjusting *α* and *β* parameters.

**Figure 12:**
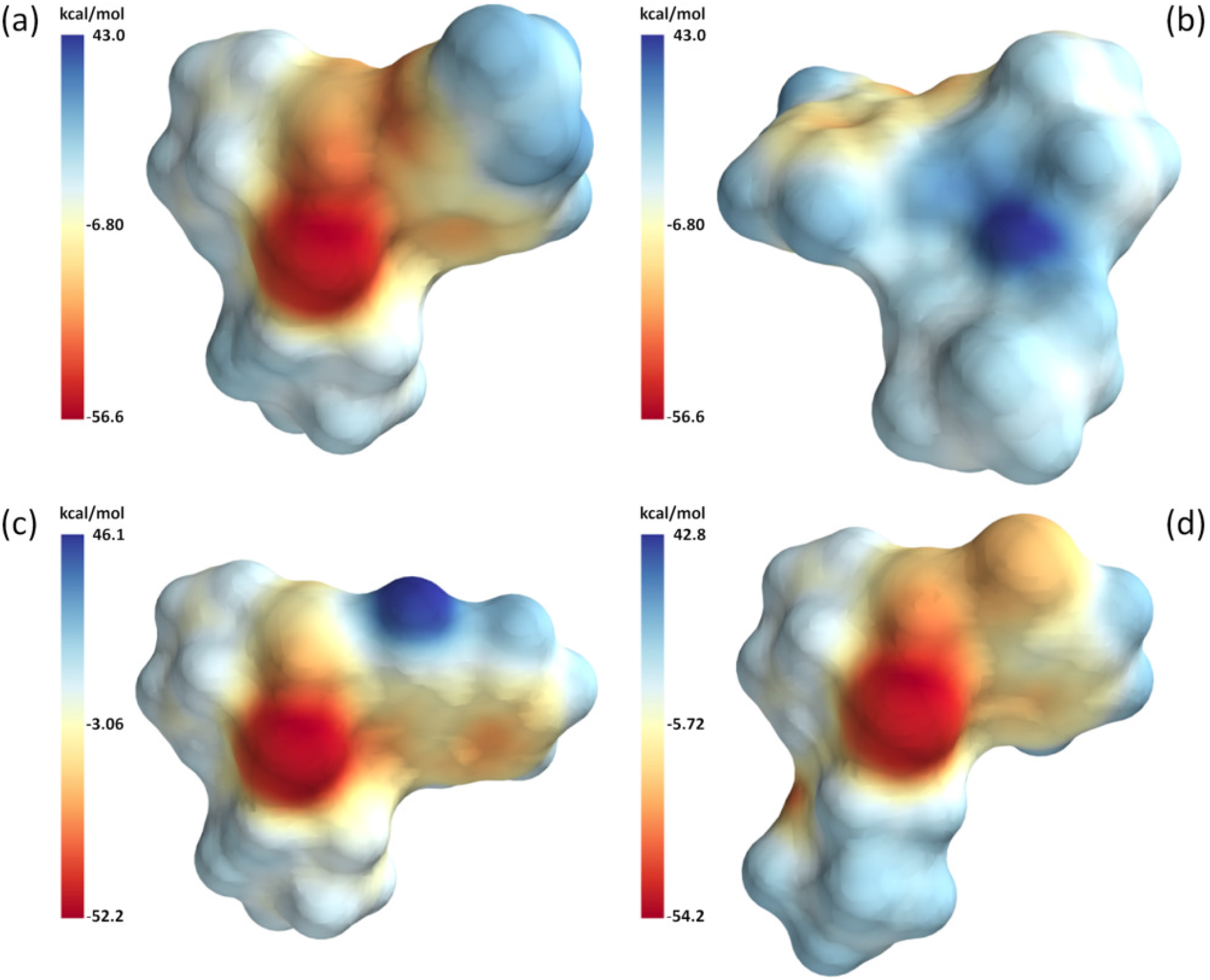
Surface Electrostatic Potential for the query molecule ZINC67007472 (a and b) and selected molecules ZINC42901627 (c) and ZINC43833531 (d) based on searches prioritizing shape or electrostatics, respectively.

Large high-dimensional datasets describing the molecular vicinity can be overwhelming for human analysis but are suitable for Machine and Deep Learning algorithms running in modern hardware. It is also possible to analyse the network graph formed by selecting pairs of *α* and *β* parameters and applying a suitable cut-off threshold to the metric. The topology of this network can yield a wealth of information about the chemical composition of the chemical space around any query molecule. If the query molecule is bound to a protein pocket, we hypothesise that using graph theory methods, it should be possible to rapidly cluster possible binders with custom selected properties. The result would be a tractable list of candidates that can be readily tested for function *in vitro*.

## 5 Conclusion

With Lira, we demonstrate that spherical harmonics can be used as the basis of an accurate and efficient platform for the search, selection and comparison of molecules, for both shape and electrostatics, given an input query. This approach can be employed to compare large numbers of molecules efficiently, establishing a robust and compact method for the description and comparison of molecular shape and electrostatic potential, that is, in addition, suitable for Machine and Deep Learning applications. We expect our package can be readily integrated with other structural software to create signatures directly based on the pocket characteristics of protein targets, thus accelerating novel lead identification. In addition, ongoing medicinal chemistry campaigns can benefit from applying Lira as the starting point for AI-guided drug development and lead maturation. The Lira code is provided with several tutorials and examples for queries using ZINC molecules.

## Acknowledgements

RWM would like to thanks Adrian Schreyer, Tom Blundell and Michele Vendruscolo for the helpful discussions about the future of rational drug design.

## Funding

This work has been supported by Fundação de Apoio à Pesquisa do Estado de São Paulo (FAPESP) grants #2012/00137-3 and #2013/20929-4. VBP and RWM thank the Rega Foundation and FWO (G0H7618N). VBP also thanks KU Leuven (C14/19/102).

## Notes

https://www.pinheirolab.com

### Competing Interest Statement

The authors have declared no competing interest.

